# Does herbivore diversity increase induced phytochemical diversity? Testing the pathway co-activation hypothesis

**DOI:** 10.64898/2026.10.25.758646

**Authors:** Umesh KP, Michal Eldar-Liebreich, Kankan Datta, Rachel Davidovic-Rikanati, Sagar Pandit, Moshe Coll, Asaf Sadeh

**Affiliations:** Dep. of Agroecology and Natural Resources, Newe Ya’ar Research Center, Agricultural Research Organization (Volcani Institute), Israel; Dep. of Entomology, R.H. Smith Faculty of Agriculture, Food & Environment, The Hebrew University of Jerusalem, Israel; Department of Biology, Indian Institute of Science Education and Research, Pune, India; Specialized Metabolism Unit, Newe Ya’ar Research Center, Agricultural Research Organization (Volcani Institute), Israel

**Keywords:** Multi-species herbivory, Induced defenses, Phytochemical diversity, Biosynthetic pathways, Phytohormone crosstalk, Ecological metabolomics

## Abstract

- The mechanistic link between multiple herbivory and phytochemical diversity as an emergent plant trait is poorly understood. Because alternative, interacting biosynthetic pathways enable high specificity of induced plant responses to different herbivores, we hypothesized that multi-species herbivory increases phytochemical diversity by simultaneously activating mutually-antagonistic response pathways.
- We tested this hypothesis using a dual-herbivory experiment with phloem-feeding aphids and leaf-chewing caterpillars on cabbage seedlings, combined with untargeted metabolomics. To isolate plant responses to herbivore identities from responses to damage levels, we equalized herbivory pressures across treatments. The underlying mechanism was tested using similar applications of the respective defense phytohormones.
- Phytochemical richness increased under dual-herbivory relative to single-species herbivory. Phytochemical evenness (^2^*D* diversity) increased from caterpillar-only to dual-herbivory. Phytohormone applications produced similar effects, supporting the hypothesized mechanism. Analyses of glucosinolate levels and whole-metabolome shifts were consistent with prevalent, but not exclusive, antagonistic JA-SA crosstalk, explaining the increased phytochemical evenness under combined exposures.
- The demonstrated general mechanistic basis suggests a broad positive relationship between phytochemical and herbivore diversity across plant–herbivore systems. Using a theoretical simulation model, we extend our two-species hypothesis to the community level. The ecological effects of herbivore-induced plasticity in phytochemical diversity may ultimately cascade through communities and food web interactions.

## Introduction

Plants produce a highly diverse array of secondary metabolites that shape defense and mediate interactions with insect herbivores. For decades, research had focused on the bioactive functions of specific phytochemicals or classes of phytochemicals, and their abundances (Fraenkel, 1959; Mithöfer & Boland, 2012). However, recent advances in metabolomics enable scientists to examine the ecological and evolutionary roles of phytochemical diversity itself, while borrowing concepts from the field of community ecology such as diversity metrics and partitioning, and emergent system-level functions (Richards *et al*., 2015; Wetzel & Whitehead, 2020; Petrén *et al*., 2024; Hanusch *et al*., 2026; Choi *et al*., 2026). Greater chemical diversity is considered to provide plants with resistance against a wider range of enemies, because no single metabolite can protect plants from all herbivores (Whitehead *et al*., 2021). Similarly, secondary metabolite diversity may also contribute to an array of non-defensive ecological interactions (Nelson & Whitehead, 2021; Slavković & Bendahmane, 2025). Thus, phytochemical diversity is a key functional trait in plants, the effects of which can also cascade up to higher trophic levels and influence community structures (Richards *et al*., 2015; Slinn *et al*., 2018; Salazar & Marquis, 2022).

Studies on phytochemical diversity have focused primarily on its evolutionary origin and maintenance (Thon *et al*., 2024), with leading theory highlighting diffuse coevolution with diverse herbivore communities (Salazar *et al*., 2018), and the interaction diversity hypothesis (Whitehead *et al*., 2021). Large scale field studies indicate that phytochemical diversity is selected for by herbivore pressure or by abiotic factors that shape biotic interactions (Defossez *et al*., 2021). For example, surveys of rainforest tree species, and of tree communities across latitudes found that phytochemical diversity varies with herbivore pressure and specialization but not with plant phylogeny (Wang *et al*., 2023; Sun *et al*., 2024). These advances raise a complementary ecological question: for given genotypes, how does the diversity of organisms that interact with plants shape phytochemical diversity dynamically, through plasticity and induced responses?

The chemical compositions of individual plants are responsive to a multitude of environmental factors, and therefore phytochemical diversity is dynamic. Recent studies showed that the diversity of neighboring plant species alters phytochemical diversity in individual plants, due to their responses to a broader array of volatiles that are emitted by these neighbors (Medina-van Berkum *et al*., 2026). Plants also interact dynamically with insect communities, particularly herbivores, and their responses can be highly specific to the taxonomic identity, feeding traits, and ontogenetic stage of different herbivores (De Moraes *et al*., 1998; Agrawal, 2000; Erb *et al*., 2012; Clavijo McCormick *et al*., 2012; Kant & Schuurink, 2021). Plasticity in plant responses to herbivore attacks is mediated by perception factors, broadly categorized as herbivore- and damage-associated molecular patterns (Erb *et al*., 2012; Karban, 2020; Monson *et al*., 2022) that activate varying combinations of phytohormones and other regulatory factors (Walling, 2000; Verma *et al*., 2025), which in turn selectively trigger specific biosynthetic pathways from a larger potential metabolic network (Mithöfer & Boland, 2012; Monson *et al*., 2022; Thon *et al*., 2024). Within this branching metabolic architecture, the central and most-studied, but not exclusive bifurcation is between the jasmonic acid (JA) pathway and the salicylic acid (SA) pathway, generally activated by leaf-chewing and sap-sucking herbivores, respectively (Walling, 2000; Ali & Agrawal, 2012; Jiao *et al*., 2024). Activating such alternative pathways ultimately leads to the induction of different suites of compounds in the plant.

When attacked by multiple herbivores, plants are challenged to activate different biochemical pathways simultaneously. Crosstalk mechanisms between biochemical pathways have been studied extensively, particularly between the JA and SA pathways. Although synergistic effects have been documented in some cases (Mur *et al*., 2006; Thaler *et al*., 2012; Ullah *et al*., 2022), interactions between biosynthetic pathways generally tend to be antagonistic, in the sense that the induction of one pathway downregulates the production of various metabolites by the other pathway (Doares *et al*., 1995; Felton *et al*., 1999; Pieterse *et al*., 2012; Monson *et al*., 2022), thus allowing the prioritization and control of alternative responses. While such antagonism has been demonstrated with respect to the abundances of specific classes of compounds, its effects on overall phytochemical diversity remain poorly understood. We hypothesized that herbivory by multiple herbivore species triggers activity in multiple response pathways simultaneously, while limiting the magnitude of each specific response. As a result, it is expected to increase metabolite richness while promoting a more even distribution of compound abundances [Fig. 1]. Alternatively, antagonistic crosstalk may lead to complete dominance of one biosynthetic pathway over the other, resulting in a phytochemical composition and diversity comparable between single- and multiple-species herbivory. However, the latter outcome is likely to occur only if the damage inflicted by (or expected from) one herbivore species is far greater than that of the other, therefore leading to a fully prioritized or canalized, rather than integrated response (de Bobadilla *et al*., 2022).

**Figure 1.**
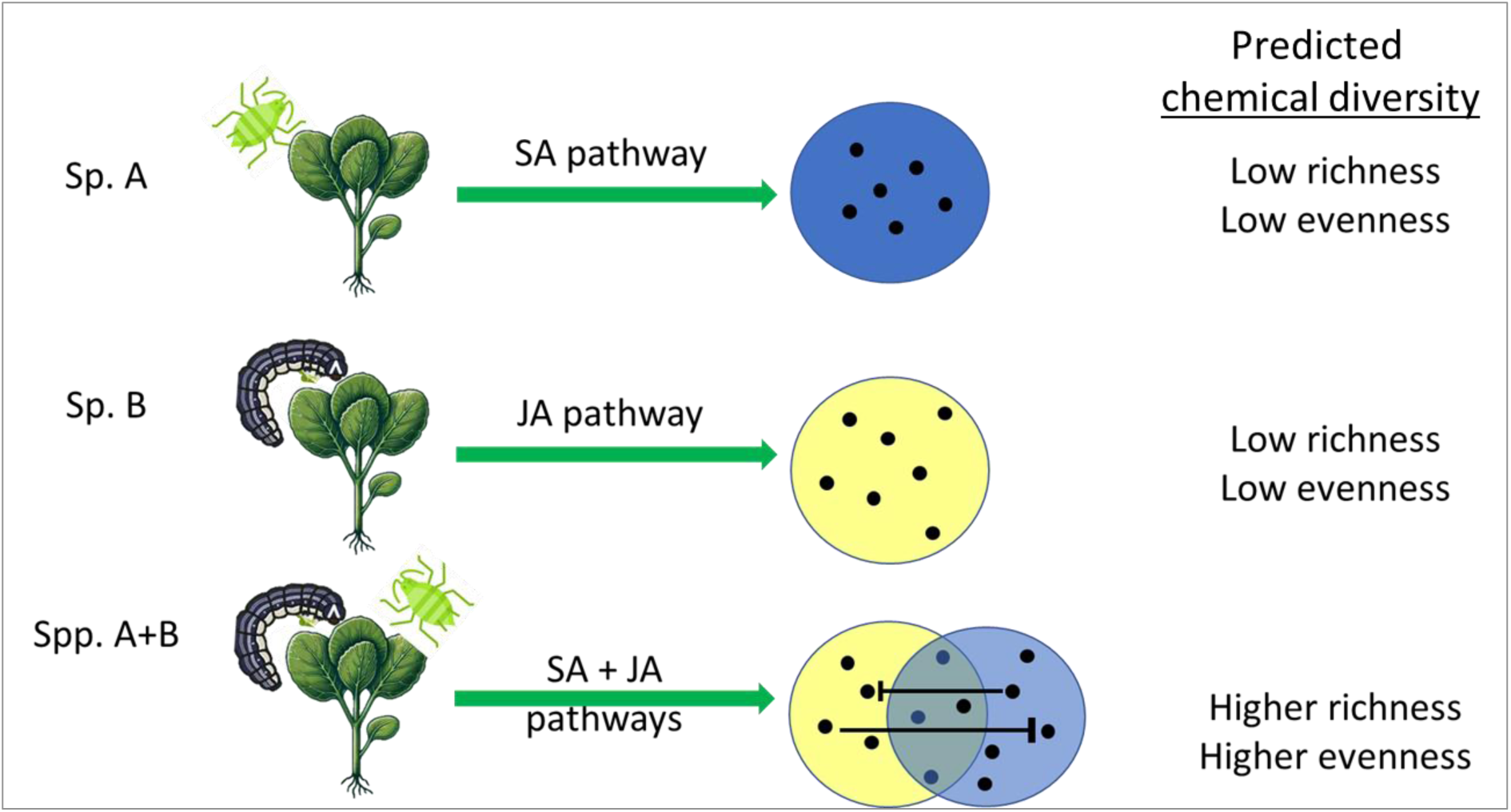
The hypothesized effect of multi-species herbivory on phytochemical diversity. Phytochemical richness increases under multiple herbivory (e.g., by aphids and caterpillars) as a result of the simultaneous activation of different biosynthetic pathways (e.g., Salicylic acid and Jasmonic acid pathways), each with its associated suite of expressed phytochemicals (depicted as Venn sets). Generally antagonistic crosstalk interactions between these pathways reduce the abundances of various product metabolites (blunt arrows), leading to greater phytochemical evenness. Thus, multiple herbivory is predicted to result in greater phytochemical diversity.

To test this hypothesis, we employed a dual-species herbivory experiment with cabbage seedlings (*Brassica oleracea*), using two generalist herbivores with different feeding modes: a sap-feeding aphid (*Myzus persicae)* and a leaf-chewing moth caterpillar (*Spodoptera littoralis).* Plants also respond to the intensity of herbivore-inflicted damage (Underwood, 2000; Choi *et al*., 2026). To isolate plant responses to herbivore species identity and diversity from their responses to varying herbivory pressure (Ali & Agrawal, 2012), we used functionally-equivalent herbivore loads (Sadeh, unpublished). To verify the underlying mechanism, we carried out a similar induction experiment using foliar application of phytohormones to artificially elicit plant responses similar to those of caterpillar herbivory (using methyl jasmonate; MeJA), aphid herbivory (using salicylic acid; SA) and dual herbivory (using combined MeJA and SA). An untargeted metabolomics approach was used to quantify the resulting richness and evenness of putative phytochemicals in the foliage. To verify the occurrence of antagonistic JA-SA crosstalk in the phytohormone application experiment, we also used a semi-targeted analysis to measure the abundances of glucosinolates, the primary group of defense metabolites in Brassicaceae. Based on the experimental results, we developed a simple simulation model to assess how broader diversity patterns of herbivore communities may affect phytochemical diversity through multiple interacting induced response pathways.

## Materials and Methods

### Dual herbivory experiment

Cabbage seedlings (*Brassica oleracea* var. capitata, cultivar Pruktor F1) were transplanted at the 3-4 true leave stage to 2.5 L containers with potting mix (Green 90, EvenAri Green Ltd.). As a phloem-feeding herbivore, we used *Myzus persicae* (Hemiptera: Aphididae), which were reared in the greenhouse at 23°C (± 2°C range) under a 16 h light: 8 h dark cycle on caged cabbage plants. As a leaf-chewing herbivore, we used *Spodoptera littoralis* larvae (Lepidoptera: Noctuidae), which were reared in the lab on *Ricinus communis* leaves up to the third instar stage, and then fed on cabbage leaves for three days before being introduced onto the experimental plants. Both insect species are highly polyphagous, and known to sometimes feed on cabbage plants in the field (Cao *et al*., 2018 ; local pest scouts, pers. com.; Mohamed *et al*., 2019).

The experiment was conducted in a greenhouse, (18 ± 2°C temperature) using 76 potted seedlings placed individually in plastic cages with 50-mesh screened windows for ventilation. On the sixth day after potting, before inoculation by herbivores, overhead photographs of each plant were taken, and the total green area was calculated by image processing, using modified Matlab code from Lati et al. (2011). This nondestructive measure of total plant leaf area served as a proxy for vegetative mass, which we verified by cutting and weighing 20 randomly selected plants immediately after photography, confirming a strong correlation (R^2^ = 0.89). Thirty-two of the remaining seedlings were assigned randomly to four treatments, each replicated 8 times: aphid-only herbivory, caterpillar-only herbivory, dual-species herbivory, and a control group in which caged plants were not exposed to any herbivory.

To isolate the responses of the plants to the identities and diversity of herbivore species from their confounded responses to varying herbivory pressures, we used functionally equivalent insect loads, aimed to equalize damage suffered by the plants across herbivory treatments, based on prior calibration experiments that used bulk insect mass and duration (Sadeh et al., unpublished; a visualization of the implemented insect loads is presented in Fig. S1). Thus, plants were inoculated with aphids on day 6 after potting (immediately after foliar photography), and with caterpillars on day 18, ca. 2 days before the end of the experiment. In the aphid-only treatment, aphids were introduced to each plant by placing an open Eppendorf tube at the base of the stem, with a pre-weighed mass of apterous aphids of mixed stages. Aphids remaining in the tube after 24 hours were removed and weighed to calculate the net inoculation mass (29.93 ± 1.00 mg (mean±SD). In the caterpillar-only treatment, each plant was inoculated with four caterpillars at the 3^rd^ instar stage, resulting in a caterpillar load of 302.54 ± 26.65 mg (mean±SD). In addition, an empty Eppendorf tube was placed for 24 hours at the base of the stem, as was done in the aphid-only treatment. In the dual herbivory treatment, plants were inoculated with the aphid (15.51 ± 2.42 mg, mean±SD) and two caterpillars weighing 150.75 ±15.28 mg (mean±SD). To correct the caterpillar inoculation masses on five plants where one of the moth larvae had perished during the experiment, we subtracted from the total inoculation weight half of the mean weight of an individual larva. The total mass of each herbivore species in the dual herbivory treatment was approximately half of the total insect weights in the respective single-species treatment. We based this decision on a testable assumption that the combined effects of aphids and caterpillars on plant mass are additive rather than interactive. Due to constraints in caterpillar availability at the required sizes, we had to deviate from the target number of caterpillars in one of the caterpillar-only replicates, and in one of the dual-herbivore replicates. This was done while adhering to the required total caterpillar mass for each plant in these treatments, in accordance with the definition of herbivore load used in the calibration procedure (Sadeh, unpublished).

The experiment was terminated 20 days after potting, by which time the plants had 8-14 true leaves. Moth larvae were removed from all the plants. Then, the plants were harvested by cutting them at the first node and weighing the entire shoot. Up to two of the youngest, undeveloped leaves and the apex of each shoot were removed and discarded to minimize ontogenetic variation in leaf chemistry, and aphids were removed from the rest of the leaves using a soft paintbrush. The aphid removal procedure was also performed on plants that were not inoculated with aphids. The remaining leaves from each plant were removed from the stem, rinsed in deionized water to remove remaining insect material, and snap-frozen in liquid nitrogen within two minutes after shoot harvest. All leaf material collected from each plant was ground, pooled together, and stored at -80°C until metabolomics analysis.

### Hormone application experiment

To confirm the mechanism underlying the responses observed in the dual herbivory experiment, cabbage seedlings were subjected to the chemical induction treatments by topical spraying of analytical standard phytohormones (Sigma Aldrich Israel) dissolved in 0.1% Triton solution. This was done in 5 blocks with 5-day gaps between to ensure a continuous availability of plants at similar developmental stages throughout the experiment. Four induction treatments, analogous to those of the dual herbivory experiment, were implemented to plants as they reached 6–7 true leaves: control plants sprayed with triton solution only, JA-induced plants sprayed with a 250 μM methyl-JA (MeJA) solution, SA-induced plants sprayed with a 800 μM SA solution, and plants sprayed with a solution combining both 125 μM MeJA and 400 μM SA. Single hormone concentrations followed those used by Yi et al. (2016), whereas half of these concentrations were used in the combined treatment, following the linear hormonal response to single-species herbivore loads observed by Sadeh (unpublished). Every day, one or two plants per treatment were sprayed with 150 ml of the respective treatment solution inside a transparent plastic cylinder. One hour after the spraying, they were moved to mesh cages in the greenhouse. Four days after the spraying, one fully expanded leaf per induction treatment was randomly selected and snap-frozen in liquid nitrogen, until a total of eight biological replicates per treatment had been obtained within 15 days. The frozen leaves were ground and stored at −80 °C until being processed for metabolomic analyses.

### LC-ESI-QTOF-MS untargeted and semi-targeted metabolomic analyses

#### Dual herbivory experiment

A 100 mg sample of frozen, ground leaf tissue from each plant was homogenized with 1 ml 80% MeOH using 2.8 mm stainless steel beads (BeadBug prefilled tubes, Sigma/Merck), vortexed for 30 sec, and shaken for another 60 minutes on repeating the vortex every 10 min. The supernatant was collected after centrifugation (5000 g) for 5 min. One ml of each sample was then filtered through Acrodisc® Syringe Filters with GHP Membrane, 13 mm with 0.2 µm (PALL, USA) and transferred to LC vials. A total of 31 samples were analyzed, as one replicate from the caterpillar-only treatment was lost.

The analysis was carried out on an Agilent 1290 Infinity series liquid chromatograph coupled with an Agilent 1290 Infinity DAD and Agilent 6530C Quadrupole Time of Flight mass spectrometer (HPLC-ESI-QTOF-MS) (Agilent Technologies, Santa Clara, USA) at the Metabolomics Unit, Newe Ya’ar Research Center. Metabolites were separated on a Zorbax Extend-C18 Rapid Resolution HT column (2.1×50 mm, 1.8 micron; Agilent Technologies) using a water–acetonitrile gradient with formic acid, and data were acquired in negative electrospray ionization mode. The gradient elution mobile phase consisted of H_2_O with 0.1% (v/v) formic acid (eluent A) and acetonitrile containing 0.1% (v/v) formic acid (eluent B). The column was equilibrated with 2.5 % B at a flow rate of 0.3 mL/min for 2 min. Eluent B was then increased to 80 % till 10 min and maintained till 12 min, then raised to 95 % B till 14 min and restored to 5 % by 15 for re-equilibration till 16.5 min. The flow rate of the mobile phase was 0.3 ml/min and the column oven temperature was 40°C. Eluting compounds were subjected to Jet Stream electrospray ionization interface operated in negative mode with the following settings: gas temperature of 315 ͦ C with a flow of 12 L/min and nebulizer set to 35 psig, sheath gas temperature of 360 ͦ C at 12 L/min flow. VCap set to 3000 V, the fragmentor to 140 V and the skimmer to 65 V. Data were acquired with a 100-1700 mass/charge (m/z) range. The electrospray capillary voltage was 3500 V (Agilent Technologies, Santa Clara, USA).

#### Hormone application experiment

To perform metabolite profiling for the hormone-sprayed plant material, 100 mg from each sampled leaf were homogenized with 1 ml 80% MeOH using 2.8 mm stainless steel beads (BeadBug prefilled tubes, Sigma/Merck) vortexed for 30 sec and shaken for another 60 minutes on repeating the vortex every 10 min. AB Sciex X500R-QTOF (UPLC-ESI-QTOF) at the Agricultural Biotechnology and Chemical Ecology (ABCE) laboratory, IISER Pune. Metabolites were separated on Phenomenex Gemini® C18 column (50x 4.6 mm, 5μm, 110Å), using a water–acetonitrile gradient with formic acid, and data were acquired in negative electrospray ionization mode. The supernatant was collected after centrifugation (5000 g) for 5 min. One ml of each sample was then filtered through Acrodisc® Syringe Filters with GHP Membrane, 13 mm with 0.2 µm (PALL, USA) and transferred to vials for Liquid chromatography/time-of-flight/mass spectrometry (LC-TOF-MS) analysis. For the UPLC-ESI-QTOF analysis, 10μl samples were separated on Phenomenex Gemini® C18 column (50x 4.6 mm, 5μm, 110Å), using AB Sciex X500R-QTOF (UPLC-ESI-QTOF) at the Agricultural Biotechnology and Chemical Ecology (ABCE) laboratory, IISER Pune. The gradient elution mobile phase consisted of H 2 O with 0.1% (v/v) formic acid (eluent A) and acetonitrile containing 0.1% (v/v) formic acid (eluent B). The column was equilibrated with 2.5 % B at a flow rate of 0.3 mL/min for 2 min. Eluent B was then increased to 80 % till 16 min and maintained till 18 min, then raised to 95 % B till 20 min, and maintained till 23 min and restored to 5 % by 25 for re-equilibration till 30 min. The flow rate of the mobile phase was 0.3 ml/min, and the column oven temperature was 40°C. Eluting compounds were subjected to Jet Stream electrospray ionization interface operated in negative mode with 5000V spray voltage, 400°C curtain gas temperature, and 10V collision energy. Data were acquired with a 100-1000 mass/charge (m/z) range.

To perform semi-targeted analysis of glucosinolates in the hormonally induced plant material, an additional aliquot of 100 mg of frozen, ground material from each sampled leaf was extracted with 1.0 mL 80% MeOH spiked with glucotropaeolin internal standard (1 µg mL⁻¹). Tubes were immediately incubated at 75 °C for 10 min to inactivate myrosinase (Doheny-Adams *et al*., 2017). The material was then homogenized in 80% MeOH using 2.8 mm stainless steel beads (BeadBug prefilled tubes, Sigma/Merck), vortexed for 30 sec and shaken for another 60 minutes on repeating the vortex every 10 min. The supernatant was collected after centrifugation (5000 g) for 5 min. One ml of each sample was then filtered through Acrodisc® Syringe Filters with GHP Membrane, 13 mm with 0.2 µm (PALL, USA) and transferred to LC vials. For LC–TOF–MS, 10 μl samples were injected onto a Phenomenex Gemini® C18 column (50x 4.6 mm, 5μm, 110Å) and analyzed on an AB Sciex X500R-QTOF (UPLC-ESI-QTOF) at the Agricultural

Biotechnology and Chemical Ecology (ABCE) laboratory, IISER Pune. The gradient elution mobile phase consisted of LCMS grade H_2_O with 0.2% (v/v) formic acid (eluent A) and acetonitrile containing 0.2% (v/v) formic acid (eluent B). The column was equilibrated with 10 % B at 0 min. Eluent B was then increased at a flow rate of 0.5 mL/min for to 85 % till 5.5 min and then raised to 90 % B till 7.5 min, and further raised to 100 % till 10 min, and restored to 3 % by 10.5 min, and then lowered for re-equilibration till 12 min. The flow rate of the mobile phase was 0.5 ml/min and the column oven temperature was 40°C. Eluting compounds were subjected to Jet Stream electrospray ionization interface operated in negative mode with -4500V spray voltage, 400°C curtain gas temperature. MS/MS fragmentation was performed with -15 V collision energy, and masses between 200-600 mass/charge (m/z) range were scanned with TOF. The semitargeted analysis identified and quantified the peak areas of three indolic (Glucobrassicin, Neoglucobrassicin, 4-Methoxyglucobrassicin), three aliphatic (Glucoiberin, Glucoraphanin [4-Methylsulfinylbutyl glucosinolate], Glucobrassicanapin) and one aromatic glucosinolate (Sinalbin) using SCIEX Analytical software. The identities of these glucosinolates were dually confirmed using their m/z values and MS-MS patterns published in the Chemspider database (Pence & Williams, 2010). In case of glucoraphinin and glucobrassicin, analytical standards were used for the confirmation of metabolite identities. Glucotrapeolin was used as an internal standard in all samples to calculate the relative peak area of the other glucosinolates. Other potential glucosinolates whose MS-MS patterns did not match with the MS-MS patterns in the database were omitted from the analysis. Lastly, the table containing the peak area of different glucosinolates was used to compute the total levels of all identified glucosinolates.

### Data analyses

The final weight of plants was compared among treatments in the dual herbivory experiment using a general linear model with experimental treatments coded as a categorical explanatory variable, and including the total leaf area measured on day 6 of the experiment as an adjustment variable. This was followed by a Tukey’s HSD test for pairwise comparisons among treatments.

For untargeted metabolomic analyses, the raw data files obtained from the Agilent HPLC-QTOF (dual herbivory experiment) as well as the raw data obtained from AB SCIEX UPLC-QTOF (hormone application experiment) were converted to mzXML file format using ProteoWizard version 3 (Chambers *et al*., 2012). The mzXML files were uploaded to XCMS Online (Tautenhahn *et al*., 2012) to perform peak picking, retention time correction, and feature alignment using the following parameters: ‘centWave’ peak picking (signal/noise threshold = 10; ppm = 20; peak width = 5, 20); ‘density’ peak grouping (minfrac = 0.7; bw = 3; mzwid = 0.005). The resulting data table with putative metabolites was exported to R statistical computing language version 3.5.1 for further trimming by feature significance, using cutoffs of p = 0.05 for the dual herbivory experiment and p = 0.01 for the hormone application experiment, to obtain the final tables of putative metabolites for each plant. A more stringent p-value threshold was applied in the hormone application experiment because the UPLC-QTOF system, used for this dataset, exhibited higher analytical sensitivity and consequently yielded a much larger number of putative metabolites compared to the HPLC-QTOF system used in the dual herbivory experiment. In the final tables, we considered putative metabolites as present in samples in which their peaks exceeded a threshold height of 5000 (Li *et al*., 2023). For the dual-herbivory untargeted dataset, blank subtraction was performed by removing any feature observed in the procedural blanks with peak height > 1100, to minimize artifacts like background contaminants and LC column carryover between samples. For a comprehensive description of phytochemical diversity in each sample, the effective numbers of putative metabolites were quantified using three Hill orders (Hill, 1973; Wetzel & Whitehead, 2020): phytochemical richness (^0^*D* diversity) was determined as the total number of putative metabolites present in the sample; ^2^*D* phytochemical diversity (highly sensitive to the evenness of metabolite abundances) was computed as the inverse Simpson index; and ^1^*D* phytochemical diversity (balancing metabolite richness and evenness) was computed as the exponential of the Shannon entropy index. These measures were calculated using the ‘vegan’ package (Oksanen *et al*., 2026), and the values of each were compared among treatments using two-way ANOVA, using package ’car’ (Fox & Weisberg, 2019), followed by Tukey’s HSD post hoc tests. In the dual herbivory experiment, one replicate in the control group was removed from the analysis after the plant was documented as growing extremely slowly, having had the lowest total leaf area on day 6 of the experiment, and the phytochemical richness value of that plant was 2.4 SDs below the mean of the control treatment. Its removal from the analysis is conservative with respect to the diversity pattern that resulted in the experiment.

The main and combined effects of experimental factors (herbivore species or phytohormone applications) on overall metabolomic composition in each experiment were tested with factorial PERMANOVA, using `adonis2` (Oksanen *et al*., 2026), based on Euclidean distances of square-root-transformed feature intensities, and type-2 partitioning of sums of squares. Significance of treatment effects was computed over 999 permutations. Homogeneity of multivariate dispersion among treatment groups was evaluated using `betadisper`. To visualize the shifts in overall plant metabolite profiles, a general linear model with an interaction term was fit to each individual feature, after auto-scaling its intensity values by their standard deviation and centering on its mean in the Control treatment. We filtered the data for responsive features with R^2^ > 0.3. Each responsive feature was classified based on its state, as “Control-associated” if its intensities in both single-herbivore (or both single-hormone) treatments were lower or similar to that of the Control, and as “aphids-(or SA-) associated” and/or “caterpillars-(or MeJA-) associated” if their intensities in the respective treatments were significantly greater than in the Control. In addition, features with significant interaction terms were also marked as having non-additive combined effects. Then, the standardized regression coefficients of each of the non-additive features were extracted, for classification of interaction types. For example, if the expected intensity of feature *i* is *E*[*y_i_*] = *β_SA_* ⋅ *SA* + *β_MeJA_* ⋅ *MeJA* + *β_int_* ⋅ *SA* ⋅ *MeJA* (with dummy-coded SA and MeJA states), then the conditional effect of SA on that feature, depending on the presence of MeJA, is *E*[*y_i_*|*SA*, do(*MeJA* = 1)] − *E*[*y_i_*|*SA*, do(*MeJA* = 0)] = *β_SA_* + *β_int_* ⋅ *MeJA*. Thus, in the presence of MeJA, the net effect of SA on that feature shows

i. ***emergence*** if |*β_SA_*| < *τ* AND |*β_SA_* + *β_int_*| > *τ*,
ii. ***nullification*** if |*β_SA_*| > *τ* AND |*β_SA_* + *β_int_*| < *τ*,
iii. ***synergism*** if *sign*(*β_SA_*) = *sign*(*β_SA_* + *β_int_*) AND |*β_SA_* + *β_int_*| > |*β_SA_*| + *τ*
iv. ***attenuation*** if *sign*(*β_SA_*) = *sign*(*β_SA_* + *β_int_*) AND |*β_SA_* + *β_int_*| < |*β_SA_*| − *τ*, and
v. ***reversal*** if *sign*(*β_SA_*) ≠ *sign*(*β_SA_* + *β_int_*) AND |*β_SA_* + *β_int_*| > *τ*

where *τ* is a tolerance threshold set to 0.1 standard deviations. Because the true biochemical contributions of each factor to the statistical interaction term are unidentifiable, our analysis represents the maximum potential directional effect of one factor on the other, calculated assuming that the effect of the conditioning factor is fixed. All analyses were conducted in R 3.5.1 (R Core Team, 2026).

## Results

### Dual herbivory experiment

Final plant weight decreased by similar magnitudes in all three herbivory treatments [Fig 2A], compared to the control (F_3,27_ = 8.87; P < 0.0003), confirming that the implemented herbivore loads had approximately equivalent impacts on the plants.

**Figure 2.**
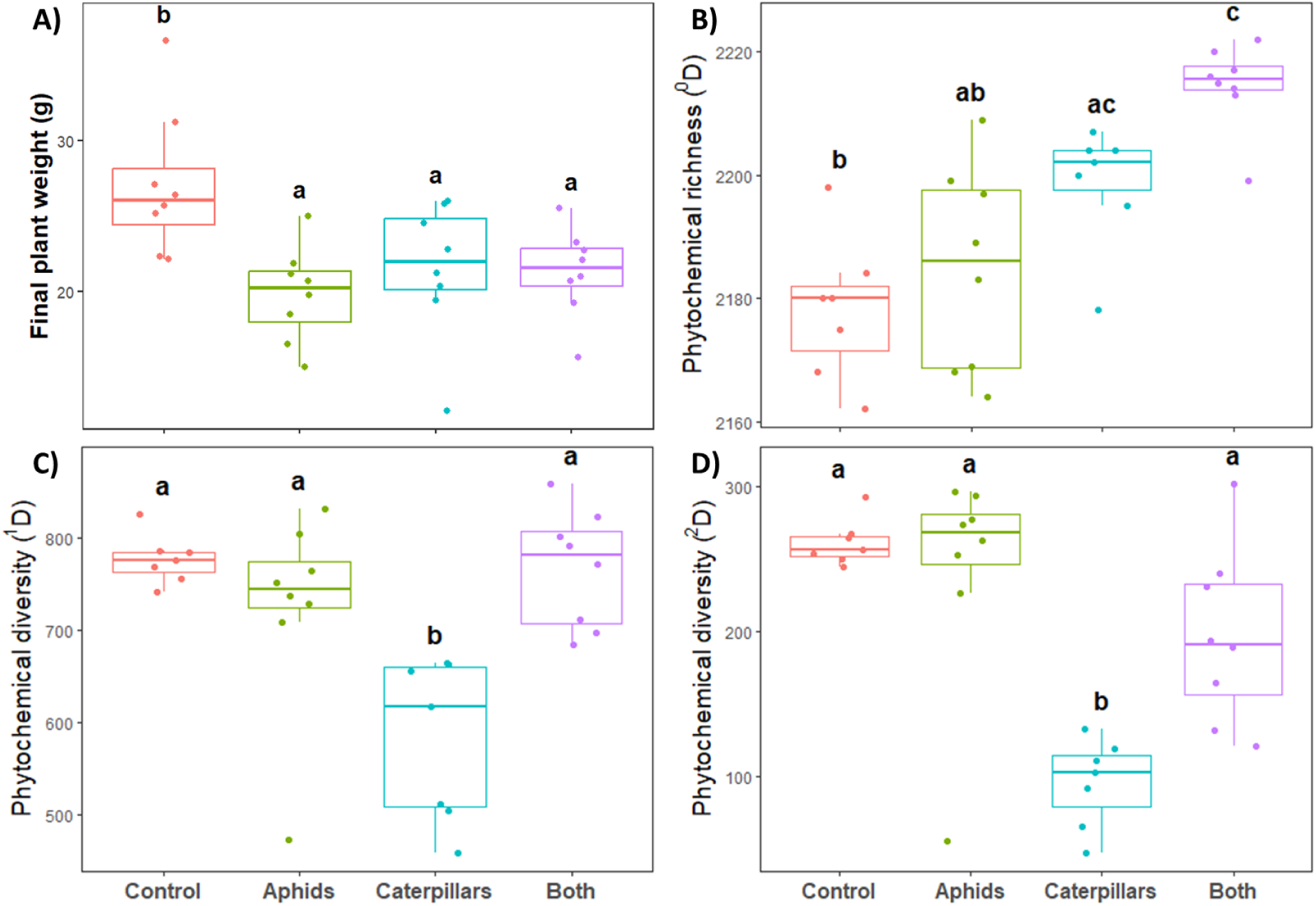
Effects of herbivory treatments on plant weight and phytochemical diversity. **A)** Similar reduction in final plant weights under all herbivory treatments, **B)** Phytochemical richness (^0^*D* diversity), **C)** ^1^*D* phytochemical diversity (exponential Shannon), and **D)** ^2^*D* phytochemical diversity (inverse Simpson, sensitive to the evenness of metabolite abundances). Differences between means with the same letter are not statistically significant (P > 0.05 in Tukey’s HSD test).

Phytochemical richness (^0^*D*) was elevated additively by aphid and caterpillar herbivory [Fig 2B] (two-way ANOVA aphids: F_1, 26_ = 6.76, P = 0.015; caterpillars: F_1, 26_ = 34.49, P < 0.001; interaction: F_1, 26_ = 1.16; P = 0.292), with lowest values in the absence of herbivory (mean 2178 putative metabolites, 95% CI: 2169 – 2187), intermediate values under single-species herbivory (mean 2185 putative metabolites, 95% CI: 2176 - 2193 for aphids; mean 2199 putative metabolites, 95% CI: 2189 - 2208 for caterpillars), and peak values under dual herbivory (mean 2214 putative metabolites, 95% CI: 2206 – 2223). ^1^*D* phytochemical diversity was affected interactively by the two herbivores [Fig 2C] (two-way ANOVA interaction: F_1, 26_ = 16.73; P < 0.001), with lowest values under caterpillar herbivory (mean 582 effective metabolites, 95% CI: 520 - 643), and higher values in the absence of herbivory (mean 777 effective metabolites, 95% CI: 715 - 838), and under aphid herbivory (mean 725 effective metabolites, 95% CI: 667 - 782). Combined herbivory resulted in ^1^*D* phytochemical diversity values similar to those of the control group despite the presence of caterpillars in this treatment (mean 767 effective metabolites, 95% CI: 710 - 825). A weaker interactive effect on phytochemical evenness (^2^*D diversity*) was observed [Fig. 2D] (two-way ANOVA interaction: F_1, 26_ = 9.16; P = 0.005), with highest values without herbivory (mean 261 effective metabolites, 95% CI: 219 - 304) and under aphid herbivory (mean 243 effective metabolites, 95% CI: 203 - 282), lowest values under caterpillar herbivory (mean 95 effective metabolites, 95% CI: 53 - 138), and intermediate values under dual herbivory (mean 197 effective metabolites, 95% CI: 157 - 236).

In a type-2 factorial PERMANOVA, the effect of caterpillars accounted for 30% and aphids accounted for 11% of the variation in the plant metabolome. A significant caterpillar × aphid interaction (F₁,₂₆ = 3.35, p = 0.022) accounted for an additional 7% of the variation. Differences among treatments in dispersion were not significant (pseudo-F_3,26_ = 2.12, P = 0.101). The metabolome’s response was characterized by 422 “control-associated” (baseline) metabolites whose intensities either decreased or non-significantly increased in response to caterpillar herbivory, and mostly decreased in response to aphid herbivory [Fig.3A]. 96 “aphids-associated” metabolites increased in response to aphids herbivory, but many of them increased also in response to caterpillar herbivory [Fig.3B]. 524 “caterpillars-associated” metabolites increased in response to caterpillar herbivory, but also showed highly variable responses to aphids. A total of 261 of these 1,015 metabolites (25.7%) demonstrated significant non-additive responses to combined herbivory. In the presence of caterpillars, the effects of aphids reversed in 83.1%, emerged in 9.2%, attenuated in 4.2%, showed synergism in 2.3%, and were nullified in 1.1% of these metabolites. Conversely, in the presence of aphids, the effects of caterpillars reversed in 23.8%, attenuated in 36.8%, increased synergistically in 31.4%, were nullified in 6.9%, and emerged in 1.1% of these metabolites (Table S1). These non-additively-responding metabolites represented 29% (154) of the caterpillar-associated metabolites, 33% (32) of the aphids-associated metabolites, and 23% (99) of the Control-associated metabolites [red lines in Fig. 3; Fig. S2A].

**Figure 3.**
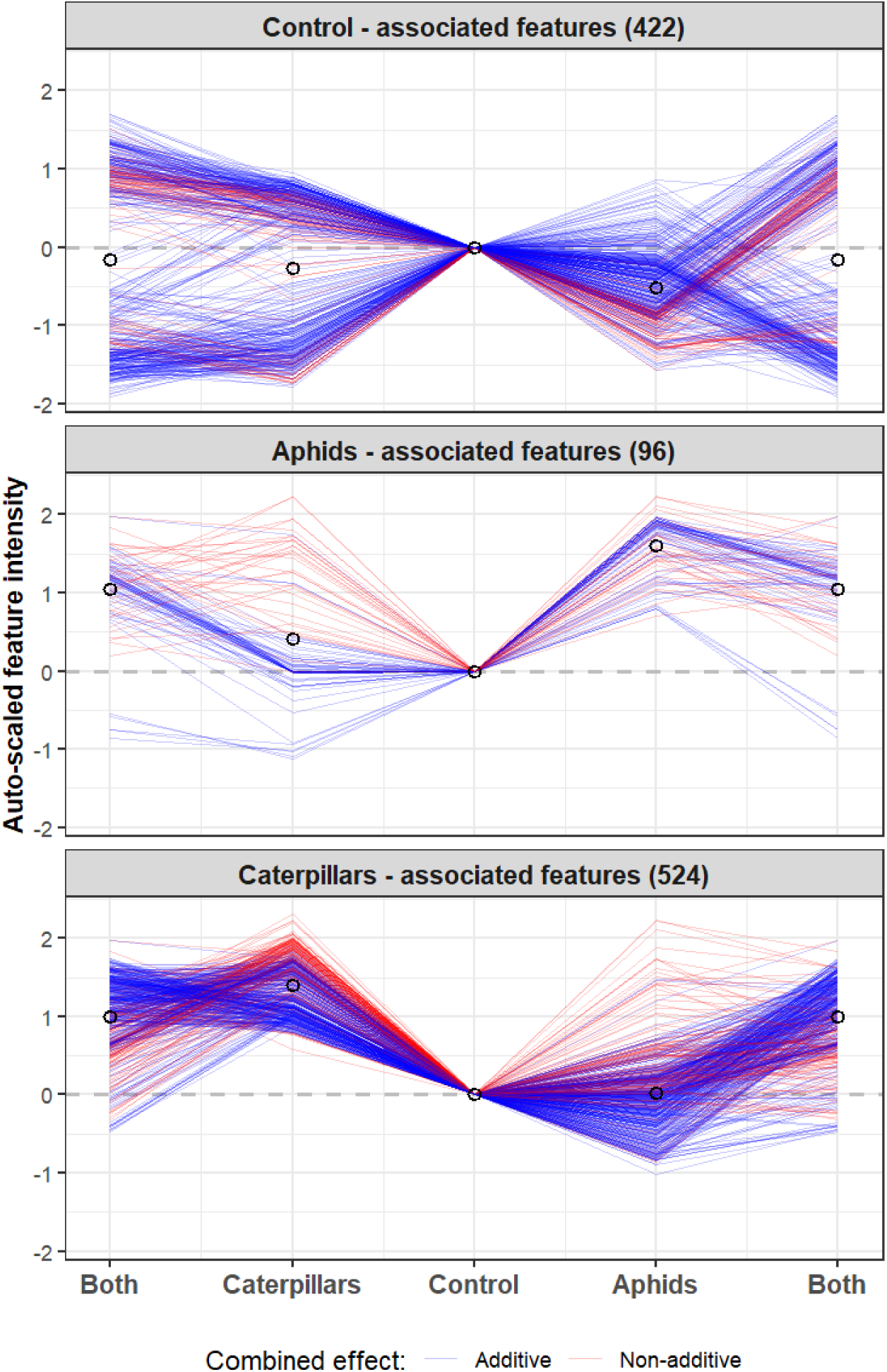
Plant metabolomic profile shifts in response to herbivory treatments. The treatments accounted for 47% (factorial PERMANOVA, P < 0.001) of the overall variation in the metabolome. The visualizations present the auto-scaled, Control mean-centered intensities of all the responsive (R^2^>0.3) features detected in the plant metabolome, grouped by their treatment associations (see Data Analyses section). Line colors indicate whether individual feature responses to herbivores are non-additive (red = significant interaction). Empty circles indicate treatment means, and dashed lines indicate the metabolites’ baseline (mean Control) values. The combined treatment is presented twice for ease of comparison with each of the single-herbivore treatments.

### Hormone application experiment

#### Untargeted analysis

Phytochemical richness (^0^*D*) was elevated interactively by SA and MeJA applications [Fig 4A] (two-way ANOVA interaction: F_1, 28_ = 7.92, P = 0.009), with peak values in plants on which both MeJA and SA were applied in combination (mean 2961 putative metabolites, 95% CI: 2872 - 3050), followed by MeJA application (mean 2709 putative metabolites, 95% CI: 2620 - 2798),

**Figure 4.**
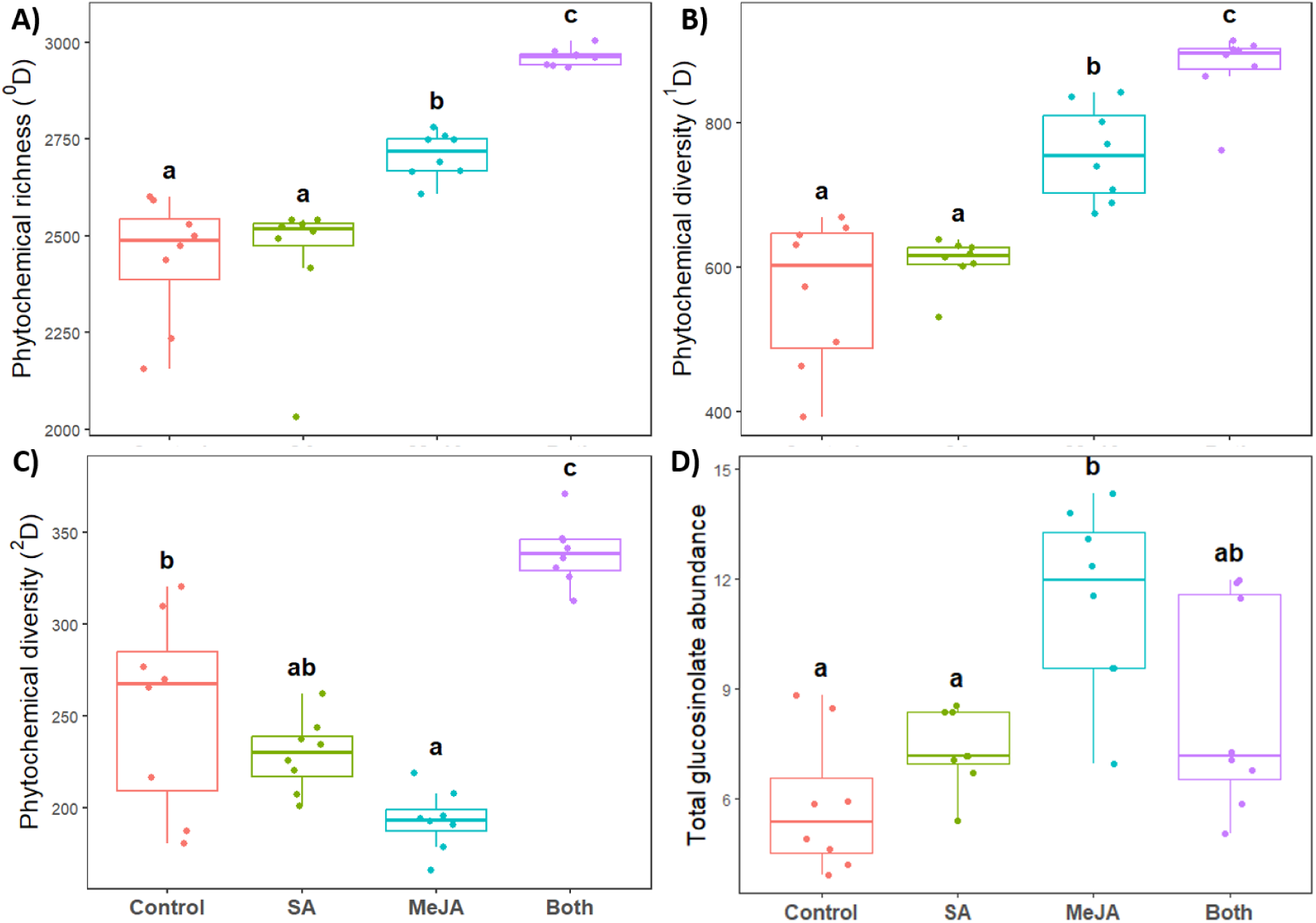
Effects of hormonal application treatments on **A)** phytochemical richness (^0^*D* diversity), **B)** ^1^*D* phytochemical diversity (exponential Shannon), **C)** ^2^*D* phytochemical diversity (inverse Simpson, sensitive to the evenness of metabolite abundances), **D)** total relative abundances of the identified glucosinolates. Differences between means with the same letter are not statistically significant (P > 0.05 in Tukey’s HSD test).

SA application (mean 2449 putative metabolites, 95% CI: 2360 - 2538), and the Control group (mean 2441 putative metabolites, 95% CI: 2352 - 2530). ^1^*D* chemical diversity followed a similar, albeit additive trend as that of phytochemical richness [Fig. 4B] (two-way ANOVA SA: F_1, 28_ = 11.38, P = 0.002; MeJA: F_1, 28_ = 92.20, P < 0.001; interaction: F_1, 28_ = 2.61, P = 0.117), with lowest values in the control group (mean 565 effective metabolites, 95% CI: 516 - 614) and the SA-treated plants (mean 607 effective metabolites, 95% CI: 558 - 657), higher values in the MeJA-treated plants (mean 757 effective metabolites, 95% CI: 708 - 806), and the highest ^1^*D* chemical diversity values were induced under combined application of SA+MeJA hormones (mean 877 effective metabolites, 95% CI: 828 - 926).

Phytochemical evenness (^2^*D* diversity) was interactively affected by the two phytohormones [Fig. 4C] (two-way ANOVA interaction: F_1, 28_ = 61.21, P < 0.001). Evenness values were similar without hormone application (mean 253 effective metabolites, 95% CI: 231 - 276) and in the SA treatment (mean 229 effective metabolites, 95% CI: 207 -251) but significantly decreased under MeJA application (mean 193 effective metabolites, 95% CI: 171 - 216). Nevertheless, the combined application of MeJA+SA yielded the greatest ^2^*D* chemical diversity values (mean 339 effective metabolites, 95% CI: 317 - 361).

In a type-2 factorial PERMANOVA, the effect of MeJA (pseudo-F_1, 28_ = 21.12, P < 0.001) accounted for 38% and the effect of SA (pseudo-F_1, 28_ = 3.42, P = 0.018) accounted for 7% of the variation in the plant metabolome. A significant SA × MeJA interaction (pseudo-F_1, 28_ = 26.51, P < 0.001) accounted for an additional 27% of the variation. Differences among treatments in dispersion were not significant (pseudo-F_1,28_ = 1.32, P = 0.305). The metabolome’s response was characterized by 1,270 “control-associated” (baseline) metabolites whose intensities mostly decreased in response to MeJA and SA applications [Fig.5A]. 549 “SA-associated” metabolites increased in response to SA application, but many of them increased also in response to MeJA application [Fig.5B]. 1,458 “MeJA-associated” metabolites increased in response to MeJA application, but most of them increased also in response to SA. A total of 1,858 of these 2,947 metabolites (63%) demonstrated significant non-additive responses to combined SA+MeJA application. In the presence of MeJA, the effects of SA reversed in 78.5%, emerged in 10.5%, attenuated in 4.8%, showed synergism in 4.0%, and were nullified in 2.2% of these metabolites. Conversely, in the presence of aphids, the effects of caterpillars reversed in 68.2%, attenuated in 24.0%, increased synergistically in 4.6%, were nullified in 2.7%, and emerged in 0.5% of these metabolites (Table S1). These non-additively responding metabolites represented 36% (671) of the MeJA-associated metabolites, 20% (379) of the SA-associated metabolites, and 59% (1,090) of the Control-associated metabolites [red line in Fig. 5; Fig. S2B].

**Figure 5.**
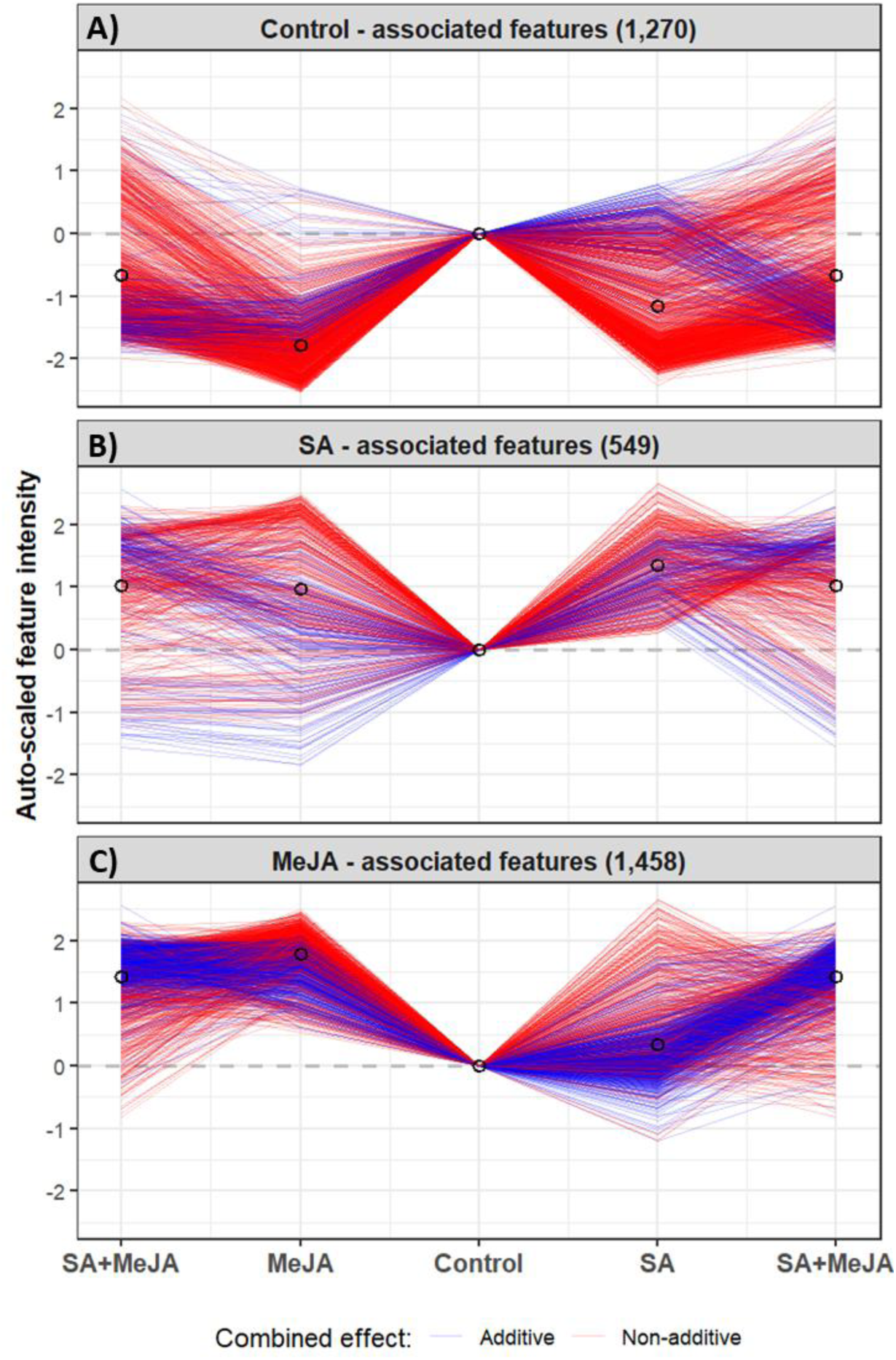
Plant metabolomic profile shifts in response to hormone treatments. The treatments accounted for 72% (factorial PERMANOVA, P < 0.001) of the overall variation in the metabolome. The visualizations present the auto-scaled, Control mean-centered intensities of all the responsive (R^2^>0.3) features detected in the plant metabolome, grouped by their treatment associations (see Data Analyses section). Line colors indicate whether individual feature responses to phytohormones are non-additive (red = significant interaction). Empty circles indicate treatment means, and dashed lines indicate the metabolites’ baseline (mean Control) values. The combined treatment is presented twice for ease of comparison with each of the single-hormone treatments.

#### Semi-targeted analysis

The total abundance of identified glucosinolates was affected interactively by SA and MeJA applications [Fig 4D] (two-way ANOVA interaction: F_1,28_ = 8.33, P = 0.007), with maximal abundance in MeJA-treated plants (mean peak area 11.41 units, 95% CI: 9.82 - 13.00), and lower abundances were detected in the other treatments (Control: mean 5.84 units, 95% CI: 4.25 - 7.44; SA: mean 7.35 units, 95% CI: 5.76 – 8.94; SA+MeJA: mean 8.43 units, 95% CI: 6.84 - 10.02). The breakdown of these responses by indolic and aliphatic glucosinolates is presented in Note S1 and Fig. S3.

## Discussion

This study demonstrates increased phytochemical richness under dual species herbivory, compared to single-species herbivory by either aphids or caterpillars, when compared under similar herbivory pressures. A similar pattern of phytochemical diversity resulted from inducing plants directly by foliar application of MeJA, SA, or their combination. In both experiments, the combination of eliciting factors resulted in additive effects on a large part of the metabolome, particularly in response to live insect herbivory, supporting our hypothesized mechanism by which multi-herbivore attack recruits a greater number of biosynthetic pathways simultaneously. Another substantial part of the metabolome responded interactively to the combination of experimental factors. These primarily involved mutually-antagonistic effects that explain the relative increase in phytochemical evenness, but also involved considerable emergent and synergistic effects. Thus, phytochemical diversity is an emergent trait that responds dynamically to varying herbivore compositions that interact with the plant, based on the structure of its crosstalk interactions network. As we discuss below, this mechanism is likely to extend beyond our model system of two herbivore species and a crucifer plant, and underlie a more general, positive relationship between herbivore diversity and the chemical diversity expressed in plants.

The interactions between the JA and SA pathways have been investigated mostly through their effects on targeted genes, compounds, or compound classes, and were found to be mostly antagonistic (Doares *et al*., 1995; Felton *et al*., 1999; Thaler *et al*., 2012; but see Davidson-Lowe *et al*., 2019), presumably because this enables plants to focus their defenses specifically towards particular herbivores that attack them (Walling, 2000; Monson *et al*., 2022; de Bobadilla *et al*., 2022). Our semi-targeted analysis of total glucosinolate levels showed a similar pattern. Although the reduced abundance of total glucosinolates in the SA+MeJA treatment could also reflect the lower MeJA concentration in the foliar spray solution, the shifts in abundances of the identified aliphatic and indolic glucosinolates suggest a mixed response to both leaf-chewers and phloem-feeders (Cipollini *et al*., 2004; Mewis *et al*., 2006; Kim *et al*., 2008; Jeschke *et al*., 2016, 2017), rather than mere dilution.

Yet, phytochemical diversity and plant function both depend on the distribution of whole-metabolome abundances (Wetzel & Whitehead, 2020). At that level, although positive interactive effects (synergism and emergence) were substantial, interactions between the factors’ effects on feature abundances were predominantly antagonistic, including many reversals (mostly of aphid effects), attenuations (mostly of caterpillar effects), and some effect nullifications. This confirms that globally, crosstalk tends to constrain the abundances of responding metabolites. Consequently, chemical evenness tends to increase in plants under multi-herbivore attack, compared to more dedicated, skewed chemical responses to single-species herbivory. Importantly, this antagonistic crosstalk does not prevent the recruitment of multiple defense pathways, thereby also increasing phytochemical richness.

A recent study by de Bobadilla et al. (2021) on plant responses to multi-species herbivory highlighted a gap between gene expression and functional defense outcomes. Our study sheds light on this question at the intermediate, metabolomic level (Peters *et al*., 2018). The distribution of interaction types in the metabolome may reflect general plant strategies for coping with multiple herbivores, such as those theorized by de Bobadilla et al. (2022), suggesting the dominance of response-integrating strategies over fully prioritized, canalized, or switching responses. Although we used functionally equivalent herbivore loads to experimentally isolate the effects of herbivore identity and diversity from herbivore pressure, plants in the field often experience imbalanced attacks, possibly leading to stronger induction of one response pathway over another and reducing phytochemical evenness. Therefore, phytochemical diversity patterns in nature are expected to emerge from the combination of herbivore functional diversity and composition, the relative pressure exerted by each herbivore group, and the interaction structure among the specific defense pathways that are triggered (Croy *et al*., 2021).

We chose to manipulate the feeding guilds or the herbivore species, and their associated JA and SA pathways, focusing on a central and well- studied branching point among alternative plant responses, in order to argue for a more general mechanism. Plant responses are known to be sensitive to additional axes of variation in herbivory other than feeding guild categories, and exhibit much greater response specificity (Tytgat *et al*., 2013; Mezzomo *et al*., 2023). This is achieved through a complex control network with many more branches than just JA and SA pathways, organized hierarchically along the biosynthetic stream (Choi *et al*., 2026). Upon exposure of cell membrane receptors to herbivore-associated molecular patterns (HAMPs) (Alborn *et al*., 1997; Schmelz *et al*., 2006), in combination with the generation of damage-associated molecular patterns (DAMPs), a cascade of signaling and regulatory factors is activated (Howe & Jander, 2008). These include, for example, calcium ion and reactive oxygen species bursts, and mitogen-activated protein kinase cascades, together with a broader hormonal network (Verma *et al*., 2025). The varying induction levels and combinations of these multi-layered factors enable a wide array of alternative fine-tuned responses, each associated with the downstream expression of different suites of phytochemicals (Choi *et al*., 2026). Consequently, if this broader control network is also dominated by antagonistic crosstalk interactions, then the diversity of herbivores in natural systems is likely to have a similar positive effect on phytochemical diversity. A stochastic simulation model of plant metabolomic response to different herbivore community compositions, assuming multiple hierarchically-branching pathways with predominantly-antagonistic crosstalk interactions (Note S2), predicts that phytochemical diversity increases asymptotically with the number of activated pathways, at a rate that depends on the functional redundancy of the attacking herbivore species (Fig. 6). Further empirical and theoretical research is needed to characterize the whole-metabolome architecture of crosstalk interactions, how it evolves in plants, and how the metabolome responds to different combinations of herbivore species, relative abundances and damage levels, and the resulting patterns of phytochemical diversity.

**Figure 6.**
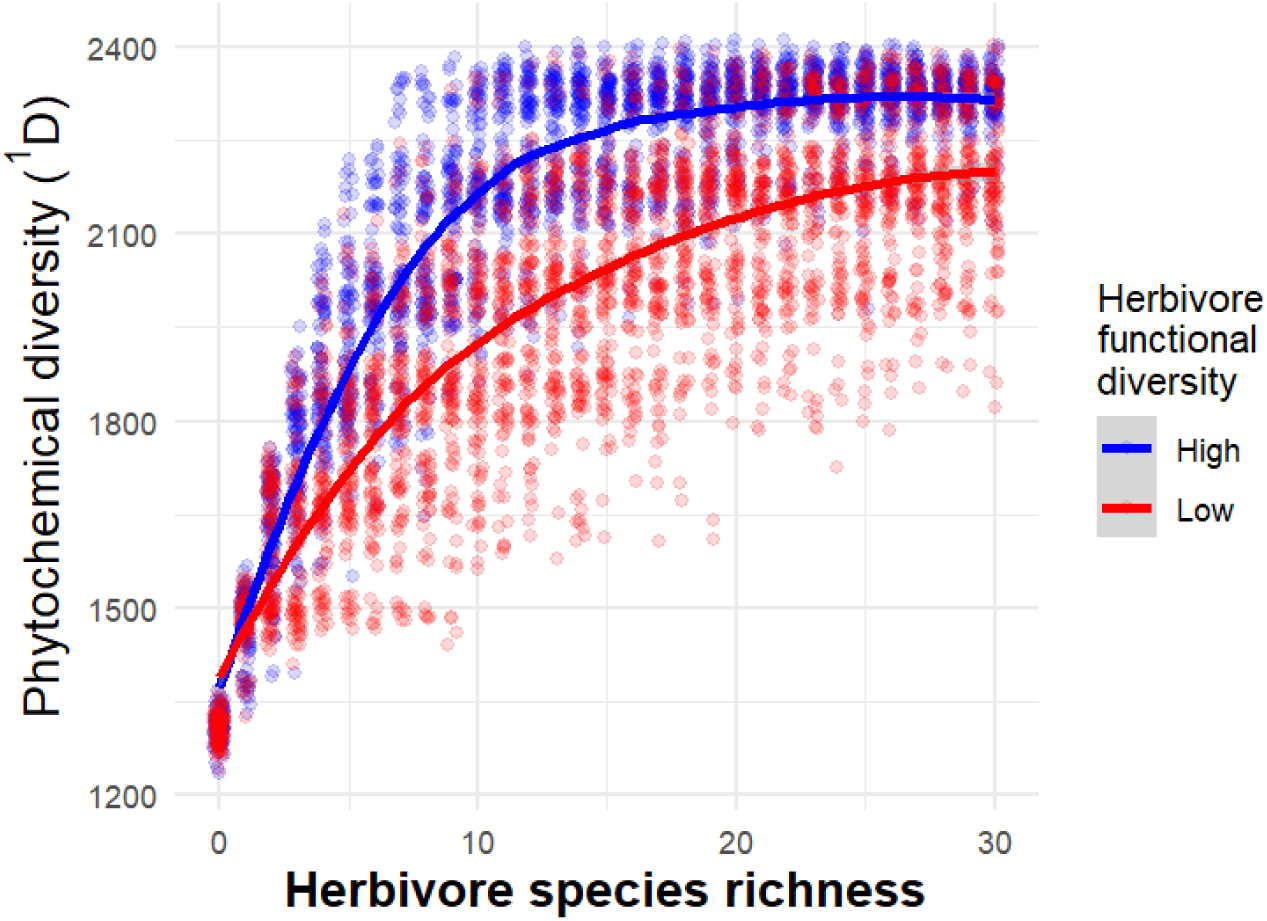
Simulated community-level relationship between herbivore assemblage diversity and within-plant phytochemical diversity. Predictions from a simulation model of metabolome crosstalk interactions based on the experimental results, in which random herbivore assemblages were drawn from species pools of high (blue) or low (red) functional diversity (e.g. diversity of feeding modes). The resulting phytochemical ^1^D diversity (exponential Shannon) values of individual plants is shown against the species richness of their simulated herbivore assemblages.

Co-variation between phytochemical diversity and herbivore diversity has been reported in natural communities (Jones & Lawton, 1991; Becerra, 2015; Richards *et al*., 2015; Salazar *et al*., 2018). These patterns have been attributed to the diversification of chemical niches for herbivores, to diffuse coevolutionary selection of plant defenses in response to the collective pressure of multiple herbivore species simultaneously, or to a positive evolutionary feedback loop between phytochemical defenses and insect specialization. We suggest that positive relationships may also emerge within individual plants at ecological time scales. Indeed, a field survey conducted on commercial lettuce (Asteraceae) found a positive association between herbivore functional diversity and phytochemical diversity in individual plants, despite spanning different genetic cultivars and agrotechnical practices (Sadeh, unpublished). Furthermore, the occurrence of this phytochemical pattern in domesticated, commercial crop plants suggests that non-domesticated plants in natural systems may generate more pronounced relationships with herbivore diversity (Liu *et al*., 2019), due to broader genomic variation that enables the induction of higher asymptotic phytochemical diversity (Fig. 6). Under growth-defense tradeoffs (Monson *et al*., 2022), herbivore diversity is expected to select for flexible plasticity and finer specificity of induced responses, as well as the ability to express multiple induction pathways more effectively, to achieve greater defense coverage. Additionally, induced chemical diversity may contribute to the evolution of constitutive phytochemical diversity if novel metabolites emerge through such multi-pathway inductions, followed by genetic assimilation (Ehrenreich & Pfennig, 2016).

In summary, our study provides mechanistic insight into how herbivore diversity can shape phytochemical diversity through multiple signaling pathways and suggests that this may be a general property of plant interactions with their insect communities. This can carry ecological implications for subsequent herbivores that may attack the induced plant (Karban, 2011; de Bobadilla *et al*., 2022). In addition to providing plants with defense against a broader spectrum of herbivores, mixtures of compounds acting through distinct physiological targets (López-Goldar *et al*., 2024) may be more challenging for individual herbivores to cope with, compared to strategies that maximize the potency of a narrow set of compounds. Thus, induced changes in phytochemical diversity are likely to feed back to herbivore population dynamics, with cascading effects on the structure and function of insect and plant communities (Glassmire *et al*., 2024). Induced phytochemical diversity may also influence the dynamics of nutritional resources in the landscape, and consequently state-dependent animal foraging patterns and health (Provenza *et al*., 1998; Pereira & Gregorini, 2022; Villalba *et al*., 2025).

## Acknowledgements

The authors thank Ruti Yonah for her logistical support, and IISER Pune for providing the infrastructure for metabolomics analyses. The project was funded by the Israeli Ministry of Science and Technology (grant # 3-16796) and Ministry of Agriculture and Food Security (grant # 20-01-0154), both awarded to AS and MC. Acquisition of the AB Sciex X500R QTOF mass spectrometer used for metabolomics analyses was funded by the Department of Science and Technology, India.

## Author Contributions

AS, MC, and UKP conceived and designed the experiments. UKP and MEL performed the experiments and collected the data. SP, KD and RR aided UKP with metabolomic analyses and pre-processing. AS and UKP conducted statistical analyses. AS developed the simulation model. All authors interpreted and discussed the results. UKP wrote the first draft of the manuscript, and all the authors contributed to the final draft; AS and MC acquired funds, administered the project and supervised the research.

## AI usage statement

No LLM was used to generate any intellectual content, including ideas, data analysis and interpretation, or conclusions. UKP used Anthropic (Claude HAIKU 4.5) to improve the clarity and readability of the manuscript text. AS consulted with Gemini Pro (3.5 Flash) for specific R syntax reminders and debugging. All the text and code suggestions were reviewed, verified and further edited by the authors, who take full responsibility for the content of the manuscript.

## Data availability

The metabolomics datasets generated during this study have been deposited in Metabolomics Workbench under project PR003366 (DOI: 10.21228/M8M00G) and will be made publicly available upon publication. The data tables and R scripts for statistical analyses are included within the article’s Supporting Information.

