## Supplemental Information for "Does herbivore diversity increase induced phytochemical diversity? Testing the pathway co-activation hypothesis"

### *New Phytologist* Supporting Information

Article acceptance date: Click here to enter a date.

The following Supporting Information is available for this article:

- **Methods S1:** Calibration of functionally-equivalent herbivore loads
- **Methods S2 – S5:** R script files for the analysis of plant weight and untargeted metabolomics in the dual herbivory experiment, and untargeted and semi-targeted metabolomics in the phytohormone application experiment
- **Fig. S1:** Insect loads implemented in the dual herbivory experiment
- **Fig. S2:** Total number of metabolites associated with each experimental factor
- **Table S1:** Classification and distribution of types of bidirectional crosstalk effects between the experimental factors on the plant metabolome
- **Tables S2 – S6:** CSV files with data for the dual herbivory experimental design and results, and the pre-processed untargeted and semi-targeted metabolomics data of the dual herbivory and phytohormone application experiments
- **Note S1:** Semi-targeted analysis of aliphatic and indolic glucosinolates
- **Fig. S3:** Effects of hormonal application treatments on total glucosinolate abundances
- **Note S2:** General description of the simulation model
- **Figure S4:** Crosstalk interaction matrix examples

**Methods S1:** *Calibration of functionally-equivalent herbivore loads*

The insect loads implemented in the dual herbivory experiment were based on a companion calibration study (Sadeh et al., unpublished). The use of those results ensured the employment of a functionally-equivalent load of *Myzus persicae* aphids and *Spodoptera littoralis* caterpillars (Fig. S1). Due to variability in aphid survival during inoculation and in their establishment on the plants, the actual aphid load was slightly higher than intended. This may be responsible for the somewhat greater (though not statistically significant) reduction of plant final weights in the aphid herbivory treatment [Fig 2A].

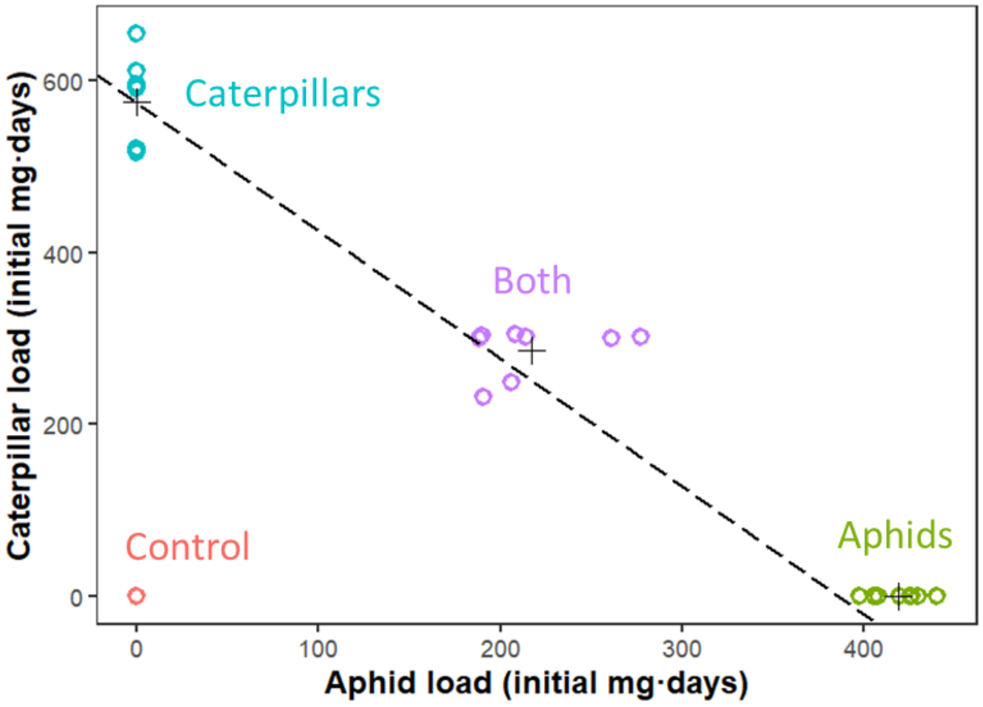


**Figure S1 |** Insect loads implemented in the dual herbivory experiment**.** Data points indicate the combinations of insect loads used for each plant, colored by treatment. Crosses indicate the means for each treatment, and the dashed line indicates the calibration slope anchored at the mean load implemented in the caterpillar-only treatment, and its linearity reflects an assumption of additive combined effects of aphids and caterpillars.

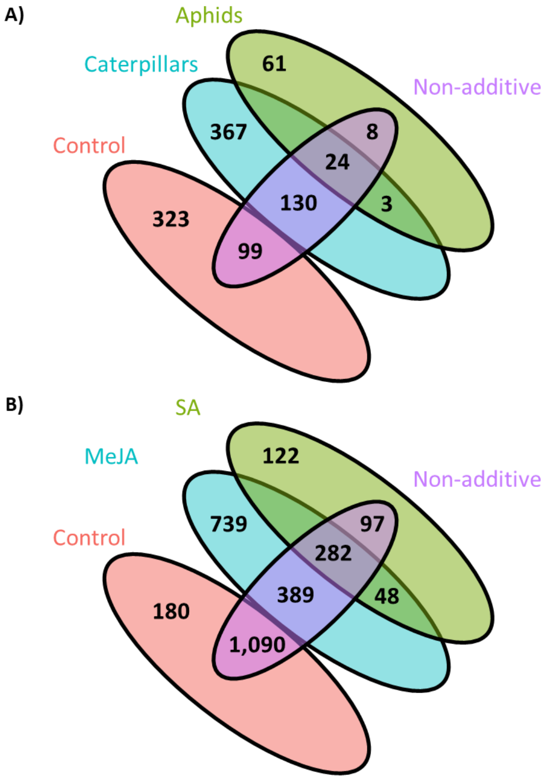


**Figure S2:** Total number of metabolites associated with each experimental factor, and those responding non-additively to the combination of factors. A) Dual herbivory experiment, B) Phytohormone application experiment.

**Table S1:** Classification and distribution of types of bidirectional cross-talk effects between the experimental factors (herbivore species or phytohormones) on the plant metabolome. Values are percent of the total number of interactively-responding metabolites.

| 261 features | Effect on caterpillars | | | | |
|---|---|---|---|---|---|
| Effect on aphids | Reversal | Nullification | Attenuation | Synergism | Emergence |
| Reversal | 19.2 | 4.2 | 28.4 | 31.0 | 0.8 |
| Nullification | 0.8 | 0.0 | 0.0 | 0.0 | 0.0 |
| Attenuation | 3.4 | 0.0 | 0.4 | 0.0 | 0.4 |
| Synergism | 0.8 | 0.0 | 1.5 | 0.0 | 0.0 |
| Emergence | 1.9 | 0.4 | 6.5 | 0.4 | ---- |

| 1,858 features | Effect on MeJA | | | | |
|---|---|---|---|---|---|
| Effect on SA | Reversal | Nullification | Attenuation | Synergism | Emergence |
| Reversal | 58.5 | 1.9 | 15.6 | 2.4 | 0.2 |
| Nullification | 0.8 | 0.1 | 0.3 | 1.1 | 0.1 |
| Attenuation | 3.8 | 0.2 | 0.2 | 0.4 | 0.2 |
| Synergism | 2.2 | 0.3 | 1.2 | 0.2 | 0.1 |
| Emergence | 3.0 | 0.2 | 6.8 | 0.5 | ---- |

***Note S1:*** *Semi-targeted analysis of aliphatic and indolic glucosinolates*

The identified aliphatic glucosinolates (Glucoiberin, Glucoraphanin [4-Methylsulfinylbutyl glucosinolate], and Glucobrassicanapin) showed overall lower abundances than indolic glucosinolates, with an increasing effect of SA application (two-way ANOVA $F_{1,28}$= 23.25, p < 0.0001), a decreasing effect pf MeJA application ($F_{1,28}$= 8.06, P = 0.008). The interaction was not statistically-significant ($F_{1,28}$= 2.21, P = 0.148), but the abundances of this compound class were similar between the SA and the SA+MeJA treatments, with lower variance in the latter [Fig. S3A].

The total abundance of the identified indolic glucosinolates (Glucobrassicin, Neoglucobrassicin, and 4-Methoxyglucobrassicin) increased strongly with MeJA application (two-way ANOVA $F_{1,28}$= 22.73, P < 0.0001), and a small, statistically non-significant increase was also observed under SA application ($F_{1,28}$= 2.44, P = 0.130), but a significant interaction ($F_{1,28}$= 10.10, P = 0.004) resulted in the majority of SA+MeJA treated plants harboring similar abundances to Control plants or those treated only with SA, although a few plants harbored as high abundances as those treated with MeJA [Fig. S3B].

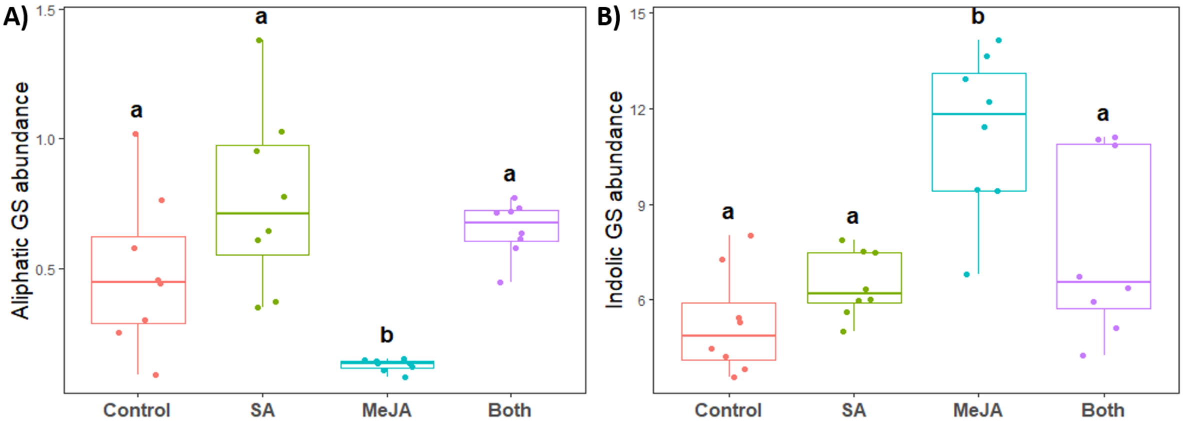


**Figure S3 | Effects of hormonal application treatments** on the total abundances of **A)** identified aliphatic glucosinolates **B)** identified indolic glucosinolates. Differences between means with the same letter are not statistically significant (P > 0.05 in Tukey's HSD test).

***Note S2:*** *General description of the simulation model*

We developed a simple stochastic simulation model of the plant metabolome and its crosstalk interaction structure. The current goal of this model was restricted to generating a basic community-level hypothesis for the effect of herbivore diversity on induced phytochemical diversity. Therefore, we did not attempt to model any of the numerous signal integration mechanisms or biosynthetic pathways studied in depth in the plant molecular biology and chemical ecology literature. Instead, we focused on the most general architectural characteristics of plant response mechanisms, as discussed in the main text. Specifically, we modeled random abundances of thousands of constitutive metabolites, defense response metabolites, and stochastically induced environmental metabolites, while imposing (i) suppressive costs of induction on primary metabolism, (ii) allocation of induction investment among main response pathways, activated based on the functional composition of simulated herbivore communities, and (iii) random, primarily negative pairwise crosstalk interactions among induced defense metabolites from different pathways. The R code for the simulation model is available upon request from the authors, and will be published in a future, more elaborated theoretical manuscript.

The model generates herbivore communities, defined by their species richness (ranging from 0 to $H = 30$) and their functional diversity. We defined $f = 6$ herbivore functional groups as groups of species that induce different primary biosynthetic pathways in the plant, as a result of their primary traits, such as feeding mode and plant organ that they attack. Herbivore communities are generated with high functional diversity by drawing species randomly from a uniform probability distribution over these functional groups, whereas low functional diversity communities are drawn from a skewed probability distribution, and are therefore dominated by species from one or a couple of functional groups. We simulated 100 cases per combination of herbivore functional diversity level and species richness values, for a total of 6,200 simulations.

In each simulation, the model focuses on the metabolomic phenotype of an individual plant. We assume the plant is attacked by the full assemblage of herbivore species occurring in its community simultaneously and at equal intensities, such that the damage caused to it by each of the attacking species is similar. Therefore, the plant responses are assumed to be shaped only by the combinations of herbivore species identities, with no variation in herbivore intensities across assemblages.

*The plant metabolome's basic structure*

The plant metabolome consists of 1,000 baseline metabolites that are always present but at variable abundances, representing products of primary metabolism and constitutive defenses. The plant is also capable of producing

2,000 environmentally-induced metabolites, irrespective of herbivory, of which a random subset of 1,000 (mean, ±50 SD) is expressed. In addition, we assumed that a primary induced defense pathway is activated by each of the herbivore functional groups upon attack by any species of that group, producing a core set of 300 specific metabolites. Each additional species from the same functional group triggers sub-branches of the primary pathway, producing additional 5 unique compounds, to a maximum of 30 species-specific compounds in each functional group. Thus, the plant can produce a maximum total of 1,980 defense-related metabolites.

The basic abundances of compounds in each of these plant's metabolite groups are drawn from lognormal distributions due to the multiplicative nature of biochemical synthesis. Their distributional parameter values (mean, SD) on the log scale are $A \sim LogN(2.5, 0.4)$ for the baseline metabolites, and $E \sim LogN(0.3, 1.2)$ for the environmentally-induced metabolites. When a single response pathway $i$ is activated, its induced metabolites spike with abundances following $B_i \sim LogN(1.5, 1.2)$.

*Crosstalk interaction structure*

Crosstalk interactions between metabolites are defined by two unidirectional pairwise interaction matrices, $M$ and $N$. Each cell in these matrices indicates the sign and magnitude of modification of an affected compound by the presence of an affecting compound, if it is expressed. We considered only inducible defense-associated metabolites as potential affecting metabolites, exerting both positive and negative effects on other inducible defense metabolites (matrix $N$), and non-positive effects on baseline metabolites (matrix $M$), both with randomly-distributed 30% sparsity. In $N$, we set the interaction values between metabolites of the same functional group to zero. Nonzero values were drawn randomly from a positive half-Normal distribution, with mean = 0 and SD = 1. Finally, we assigned negative signs to a randomly-selected proportion (70%) of the cells, while retaining their absolute values. Figure S4 presents examples of both interaction matrices.

*Cost of defense induction*

Plants that do not experience any herbivory (species richness = 0), do not express any induced defense metabolites, and therefore their constitutive metabolite expression remains unchanged. Upon exposure to any herbivore functional group (regardless of species richness within that group), induction of the respective defense pathway incurs a metabolic cost, $c$. This cost can be considered as a fraction of stored plant resources that are reallocated to defense, resulting in a proportional reduction in the abundances of baseline metabolites. The same proportional cost of the plant's remaining stored resources is incurred multiplicatively by each additional pathway induced by different herbivore groups. Thus, the overall cost of defense is

$$C(n) = S \cdot [1 - (1 - c)^n]$$

where $n$ is the functional richness of herbivores attacking the plant, and $S$ is the fraction of the plant's resource pool that is stored as reserves (the complementary fraction being resources dedicated to plant growth). This cost is distributed among the constitutive metabolites according to their interactions with the induced pathways. Letting the binary vector $x = (x_1, x_2, \ldots)$ indicate the set of metabolites produced by the plant given its interactions with its simulated community, the product $d = Mx$ is the vector of cumulative raw crosstalk effects of all the induced defense metabolites on each of the baseline metabolites. The proportional suppression of baseline metabolite abundances, $A$, scales exponentially with the global cost of induction, $C(n)$, and with the mean-normalized total effect of crosstalk on each metabolite,

$$A' = A \cdot \left[ b + (1 - b) \cdot e^{-C(n) \cdot \frac{d}{\mathrm{mean}(d)}} \right]$$

with the floor parameter, $b = 0.25$, ensuring that primary metabolism never drops below a critical proportion of the baseline abundance.

*Induced defense-associated metabolite abundances*

When a single response pathway $i$ is activated by attacks of herbivore species from a single functional group, the associated set of metabolites are induced with abundances $B_i$. When multiple pathways are activated simultaneously, given our assumption of equal herbivore pressures, each pathway is allocated an equal share $p_i = C(n)/(n \cdot C(1))$ of the total defense investment, reducing its overall magnitude to $p_i \sum \mathrm{B}_i$. In addition, the relative abundances of individual metabolites associated within each induced pathway are modified by crosstalk interactions with the other active pathways. Given $x$, the metabolite activation indicator vector, $d_i = Nx$ is the vector of cumulative raw crosstalk effects on each induced metabolite of group $i$ by the metabolites of the other groups. These net raw effects are scaled through a logistic function with floor and ceiling parameters $b = 0.25$ and $u = 4$, respectively:

$$\mathrm{X}_i = B_i \cdot \left[ b + \frac{u - b}{1 + \left(\frac{u-1}{1-b}\right) e^{-d_i}} \right]$$

and normalized by their sum, $\sum \mathrm{X}_i$. Thus, the final abundances of induced metabolites in pathway $i$ become:

$$B'_i = \frac{\mathrm{X}_i}{\sum \mathrm{X}_i} \cdot p_i \sum \mathrm{B}_i$$

The final plant metabolome, resulting from its baseline primary and constitutive metabolites, defense-associated metabolites induced by the herbivore community, and stochastic environmentally-induced metabolites is $[A', B'_1, \ldots, B'_n, E]$. For each simulated plant, we computed $^1$D phytochemical diversity over its entire metabolome.

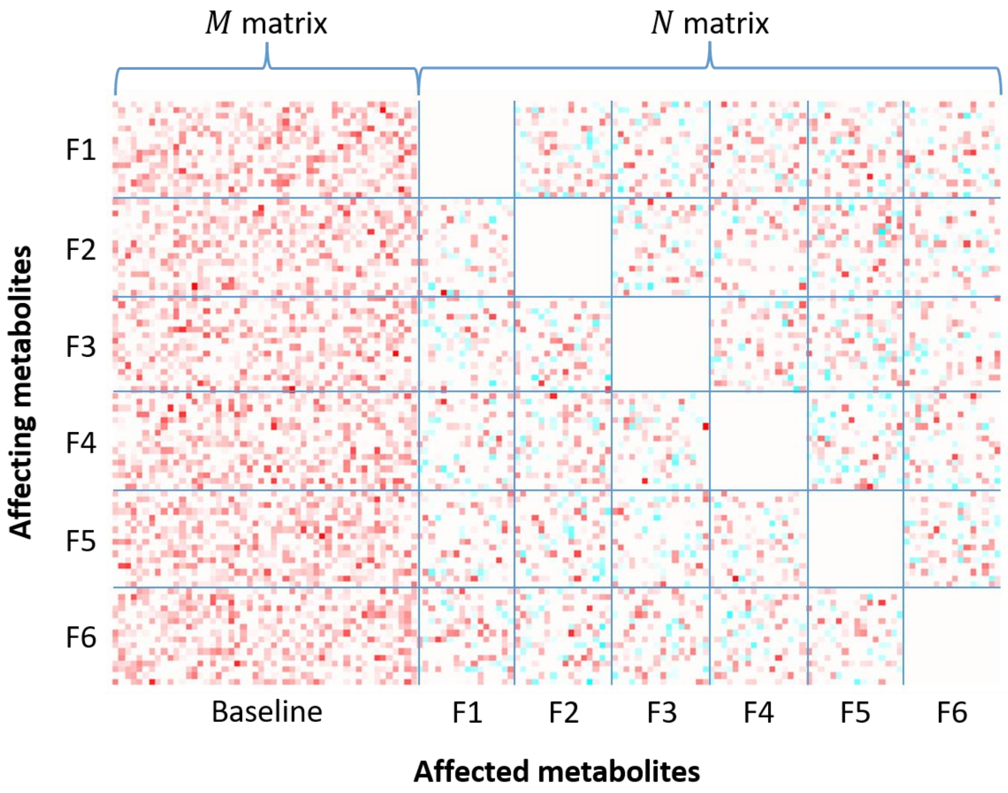


**Figure S4:** Crosstalk interaction matrix examples. Rows represent affecting, inducible metabolites (if they are activated), organized by functional-group associated induction pathway F1 through F6. Columns represent affected baseline metabolites ($M$ matrix) and inducible metabolites ($N$ matrix). Color intensity indicates the magnitude of a pairwise raw crosstalk effect (red is negative, cyan is positive, and white indicates no interaction). The number of overall metabolites presented here is reduced 20-fold for clarity.
